# Real-Time Spatiotemporal Filtering for Artifact-Free EEG during Electrical Neurostimulation

**DOI:** 10.1101/2024.11.06.622272

**Authors:** David Menrath, Joshua P. Woller, Alireza Gharabaghi

## Abstract

Combining electrical neurostimulation with electroencephalography (EEG) for adaptive neurostimulation remains challenging due to the presence of stimulation artifacts in the recorded signal. Interpretation of EEG activity concurrent with stimulation requires real-time filtering of this noisy signal. While traditional frequency domain filters can suppress activity within frequency bands, they fail to differentiate sources in situations when there is a shared frequency characteristic between brain activity and stimulation signal. Here we present a new real-time denoising approach that combines spatial filtering and dynamic filter application. The spatiotemporal filter can suppress stimulation artifacts that share frequency bands with the brain signal in real-time while preserving the full spectral power. The spatial filter is also dynamically updated to account for possible changes in artifact topography. Finally, the filter is robust to changes of stimulation intensity, frequency and duration, enabling reliable denoising performance for inclusion in brain state-based closed-loop stimulation.

## 1 Introduction

Electrical neurostimulation holds promise as a treatment for psychiatric and neurological disorders by modulating impaired neural circuits Bhattacharya et al. [2022]. To advance real-world applications, it is crucial to evolve beyond basic, preset, non-adaptive stimulation protocols. Compared to pharmacological approaches, electrical neurostimulation offers precise temporal modulation of brain activity, enabling targeted interference with specific neural processes. However, this precision remains underutilized if the stimulation is not aligned with the brain’s current state. While EEG can record brain activity with the necessary temporal resolution to facilitate adaptive stimulation, artifacts induced by electrical stimulation pose a major challenge to real-time analysis. These artifacts limit the efficacy of closed-loop neurostimulation, often restricting analysis to EEG segments recorded between stimulation bursts and omitting immediate or short-term evoked responses occuring within those bursts. To address this limitation and support closed-loop applications, removing stimulation artifacts from EEG data in real-time is a core development objective Bergmann [2018].

Common artifact attenuation strategies, such as low-pass or notch filtering, while effective, result in the irreversible loss of neural signals in the filtered frequency range. Therefore, these filters should only be employed when the stimulation frequency is outside the range of interest. Additionally, real-time applications must prevent ringing artifacts that may arise from or spike- or step-like stimulation waveforms de Cheveigné and Nelken [2019]. Beyond their spectral characteristics, many neurostimulation artifacts have consistent topographies across EEG channels, which can be extracted from channel covariances using multivariate analysis methods. While blind decomposition methods such as Independent Component Analysis (ICA) or Principal Component Analysis (PCA) can recover these artifact topographies, they are not specifically optimized for this purpose and may yield mixed components that contain both brain activity and stimulation artifacts. Conversely, Generalized Eigendecomposition (GED) Cohen [2022] as a supervised multivariate approach, can maximize any contrast of spatial patterns defined by the researcher by an appropriate choice of covariance matrices that are jointly diagonalized. In the context of neurostimulation, the occurrence of stimulation periods is known to the experimenter. Utilizing this prior knowledge by jointly diagonalizing covariances of stimulation and stimulation-free epochs facilitates the extraction of artifact topographies that efficiently capture stimulation artifacts in a single (or few) components, while removing as little neural signal as possible.

Although an artifact topography can be continuously removed by permanently applying a spatial filter (e.g. projecting the artifact topography out) throughout the entire recording, as is typical for ICs, this is not necessary if the artifact has defined onsets and endpoints. Accordingly, it may be advantageous to dynamically apply the spatial filter during active stimulation periods only.

Furthermore, alterations in the impedance of the EEG electrodes over the course of a recording session, may result in modifications of the artifact scalp topography of the stimulation artifacts, thereby necessitating an update of the spatial filter. This filter updating has been identified as a need for the denoising of EEG with concurrent neurostimulation Haslacher et al. [2021]. To perform such updating in real time, the filter calculation must have a low computation time and provide natural ordering of calculated source components to minimize the need for manual component selection.

Here, we present a framework for adaptive real-time denoising of EEG data from electrical stimulation artifacts by temporally applying a spatial filter to the incoming data stream. As this allows for an uninterrupted readout of ongoing brain activity, real-time denoising of EEG data during neurostimulation enables the development of true closed-loop adaptive stimulation protocols, which go beyond mere brain-state triggered open-loop protocols Bergmann [2018]. In this study, we used this method for cleaning EEG data from the artifacts of transcutaneous auricular vagus nerve stimulation (taVNS). It should be noted, however, that the method can be applied to remove artifacts from other electrical neurostimulation methods equally well.

## 2 Materials and Methods

### 2.1 Data

To validate our approach, we used both real human EEG data and synthetic EEG recordings from a physical head phantom model. In the human recording, the participant received taVNS while instructed to focus on a fixation cross, presented on a computer screen. TaVNS was delivered at a frequency of 25 Hz (250 μs pulse width, biphasic waveform, at 1.2 mA) over a period of 4s, followed by a 11.3 s off interval using a tVNS-R Stimulator (tVNS technologies GmbH, Erlangen, Germany). The stimulation was triggered remotely via a PC running proprietary software (tvnsmanager.exe), which communicated with the stimulator via the Bluetooth low energy protocol. The EEG data were recorded using an active EEG system with 32 channels (actiCap & BrainAmp DC, BrainProducts GmbH, Gilching, Germany). The young healthy participant provided informed consent to participate in the recording. The study was approved by the Ethics Committee of the University Hospital Tübingen (178/2023BO1).

The physical head phantom model was assembled according to published instructions Hairston and Yu [2019]. Monofrequency sinusoidal oscillations of 10 Hz with a varying spatial pattern were injected using auxiliary cables and an external sound card. A taVNS electrode was placed at the ear of the head model, and taVNS was applied at a frequency of 25 Hz (250 μs pulse width, biphasic, at 3 mA) using a tVNS-R stimulator. The EEG was recorded using actiCap slim electrodes and an actiCHamp amplifier (both Brain Products GmbH, Gilching, Germany). Both EEG recordings were sampled at 1000 Hz.

The data also included two trigger channels, one signaling 0.1 s before stimulation onset and the other 0.1 s after the end of a stimulation burst. These were used to infer the current stimulation state for the dynamic filtering.

### 2.2 Algorithm Overview

For the real-time streaming of data, we used the mne-lsl package, an interface to the Lab Streaming Layer (LSL) library which is compatible with the mne-Python functionality. In order to facilitate real-time processing, custom Python code was used.

Incoming data were obtained from the LSL stream inlet. The trigger channels were used to infer the stimulation state, i.e., whether the current sample belongs to a stimulation or to a non-stimulation period. Data from the stream were accumulated continuously into epochs of 0.5s duration, belonging either to a stimulation or non-stimulation period. From each epoch, the covariance of the EEG channels was computed, and transferred to a separate first-in-first-out queue for each stimulation state, each queue encompassing 10 such covariance matrices. As this queue is updated throughout the recording, it allows for a continuous updating of the spatial filter. The averaged covariance matrices are used to compute the spatial filter matrix using generalized eigendecomposition. This filter was then passed to a mixer to apply it dynamically contingent on the stimulation state, i.e., to blend it in only after the onset of a stimulation train was indicated from the trigger channels. This mixer was then applied to the incoming data sample, and the resulting denoised data was passed to an LSL stream outlet. A flowchart of the algorithm is illustrated in Figure 1.

**Figure 1:**
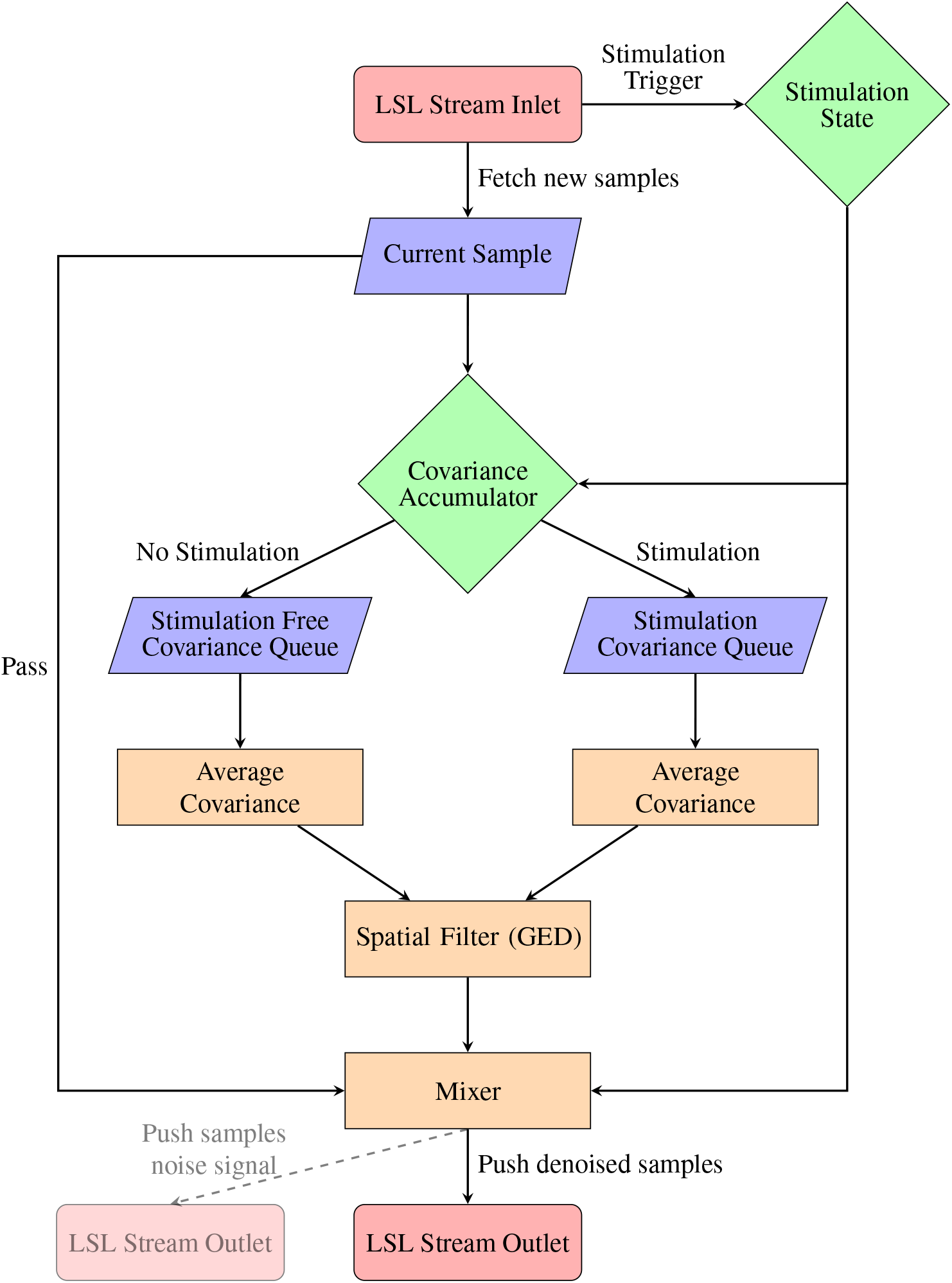
The flowchart depicts the proposed approach to spatiotemporal filtering. The data from the LSL stream inlet are assigned to a stimulation state on the basis of the accompanying trigger channels, and the corresponding covariance matrices from each stimulation state are used to compute the spatial filter. This filter is then applied dynamically to the data, contingent on the stimulation state. The cleaned data are then pushed to an LSL stream outlet for further analysis. If desired, the complementary signal (the isolated noise signal) can be passed via a second LSL stream outlet.

### 2.3 Technical Details

#### Generalized Eigendecomposition

The generalized eigenvalue problem for two matrices *A* and *B* is to find sets of scalars *λ* and non-zero vectors *x* such that:

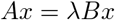

where:

- *A* and *B* are square matrices of the same dimension *n × n*,
- *λ* is a scalar (the generalized eigenvalue),
- *x* is a non-zero vector (the generalized eigenvector).

This is a joint diagonalization of the matrices A and B. For our algorithm, we reduced the generalized eigenvalue problem for symmetric matrices to two singular value decompositions, as outlined previously Melzer [2004]. This approach further may also be designed to be robust to potential rank-deficiencies of the input matrices. Eigenvalue or singular value decompositions are fast and deterministic numerical operations, having a distinct run-time advantage compared to the slower non-deterministic, iterative computation of ICA Cohen [2022]. GED can be used to construct spatial filters that enhance researcher defined contrasts, and has some applications in the denoising of EEG data during concurrent neurostimulation Haslacher et al. [2021]. For a more complete overview of the technicalities and use cases of this technique, the reader is referred to Cohen [2022].

#### Updating of the Spatial Filte

Dynamic updating of the GED is achieved by maintaining a first-in-first-out queue of covariance matrices from each stimulation condition. The contributions of these covariance matrices to the average can be adjusted by different weighting coefficients (e.g., exponential decay of weights), allowing the GED to be more sensitive to recent changes in artifact topographies, as indicated in the covariance patterns.

## 3 Results

The algorithm was efficient in removing the artifacts resulting from taVNS application. For the selected data set, the removal of two components was sufficient to subdue the stimulation artifacts in all channels (see fig. 3 for the effects on the channels most affected by the artifact). The overall spectral composition of the signal remained largely unaltered, see fig. 2. The proposed algorithm introduced only a minimal and consistent temporal delay between the acquisition of the raw data and the output of the processed data in the sub-millisecond range.

**Figure 2:**
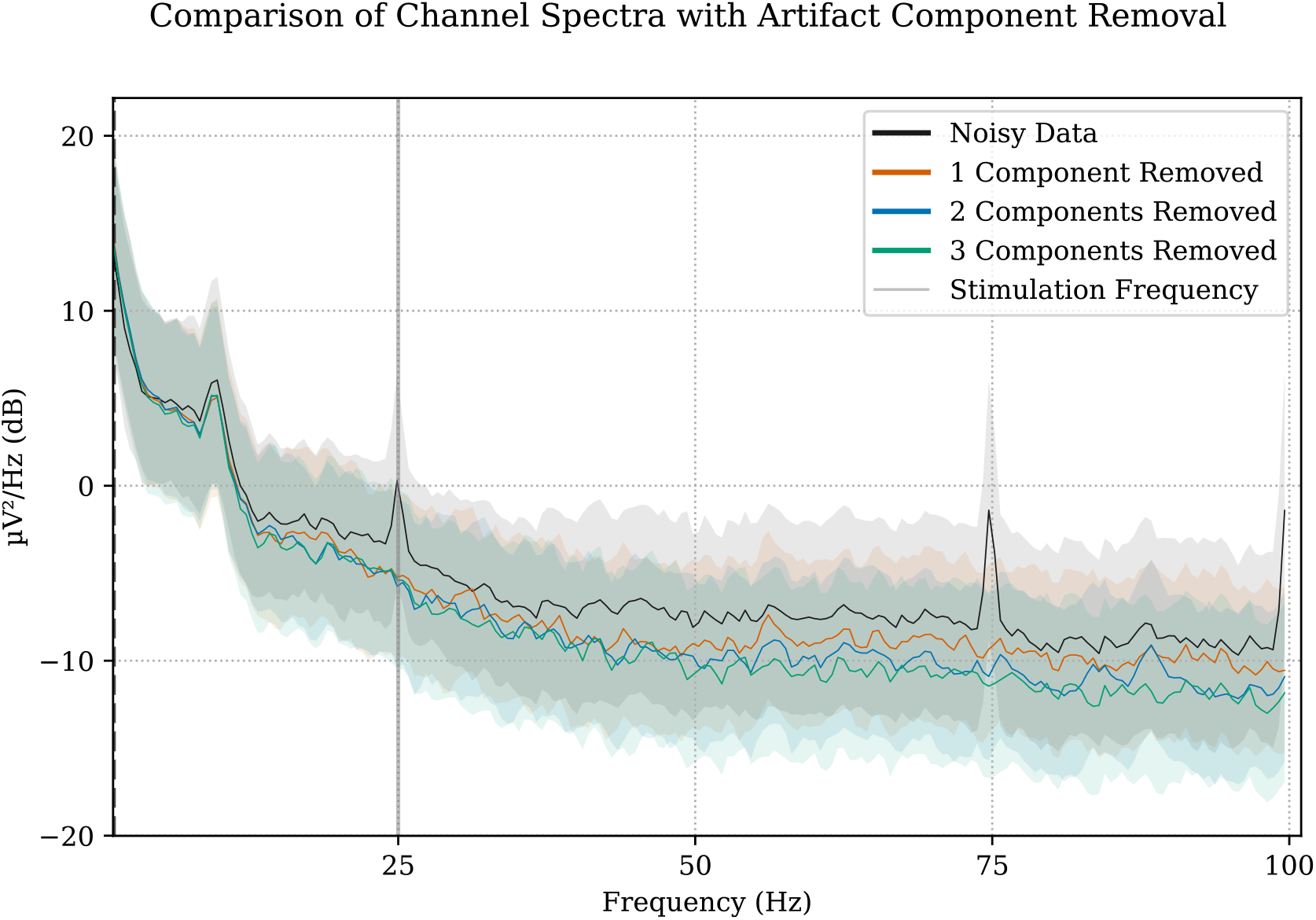
Power spectral density of the human EEG recording before and after filtering. The power spectra of the raw signal (with stimulation noise) and the filtered version after subtracting one, two or three components from the spatial filter are plotted. GED reliably removed increased power at the stimulation frequency (and its harmonies) while leaving baseline power unaltered (unlike for example notch filtering).

**Figure 3:**
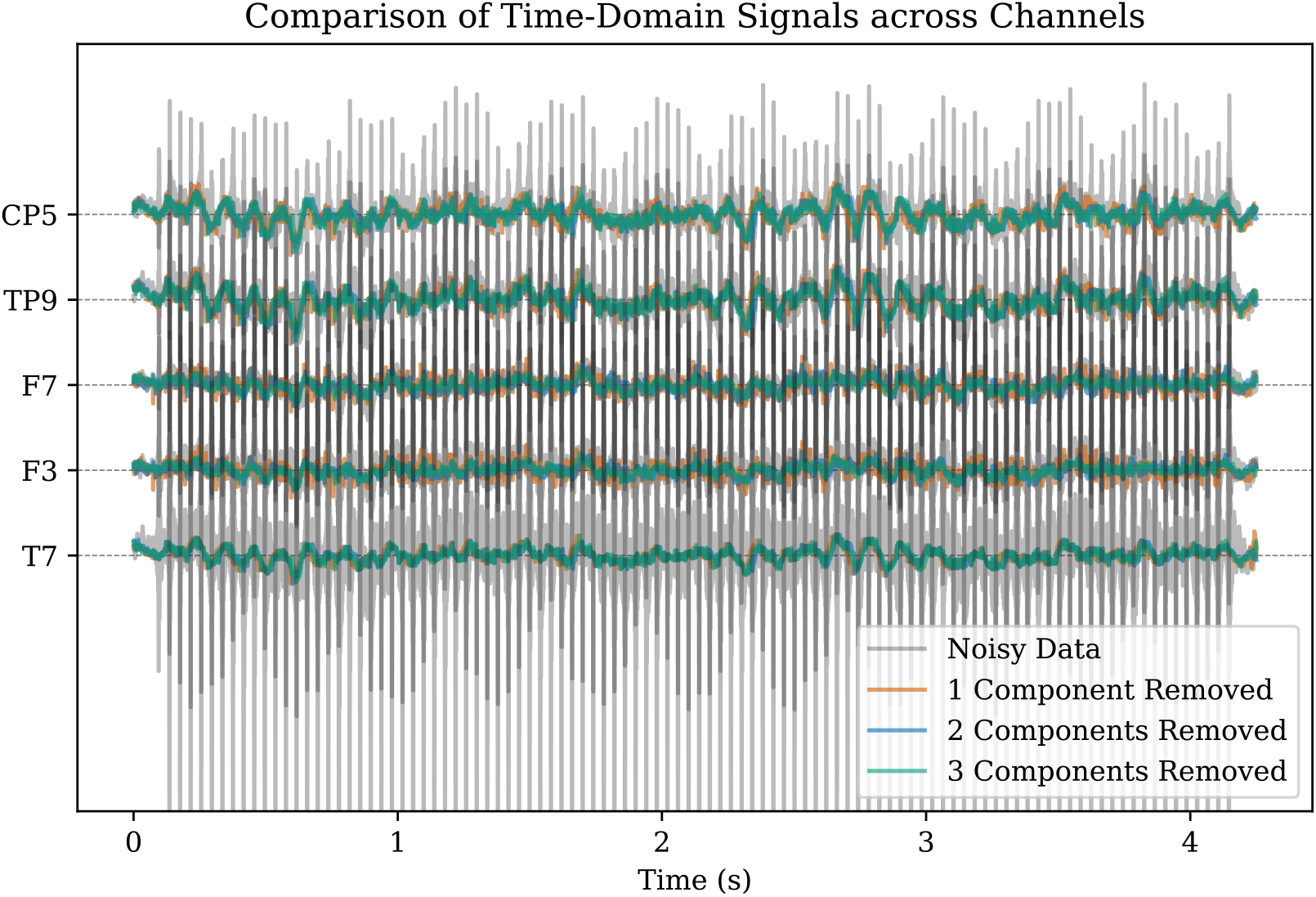
Snippet of representative EEG data collected during a taVNS burst. Colors indicate various strengths of denoising, i.e., number of components projected out. For this plot, only channels showing strong artifacts were chosen, with T7 being affected the most. While for a single component, minor spike artifacts remain, these are not visible when projecting out two or more components. A recovery of underlying slower oscillations is apparent.

## 4 Discussion

The design of the presented algorithm offers a number of improvements to the currently used denoising approaches. Using GED for the calculation of the spatial filter leverages our knowledge about the stimulation state into the filter creation, while preserving the filter’s ability to differentiate between brain signal and noise artifacts in the same frequency band. The filter calculation for GED provides substantial support for incremental updating by storing the covariances of incoming samples during normal operation. This means that the filter updating can be performed in the background, omitting the need to interrupt ongoing recording sessions for filter recalibration. This increases the reliability of the filter over longer recording sessions.

Furthermore, frequency invariance of spatial filters can greatly benefit future neurostimulation research exploring the frequency parameter space or enable subject-specificHallo, frequency tuning of neurostimulation methods.

Another benefit of spatial filtering is that the signal is decomposed into complementary components, which can be added together to reconstruct the original unfiltered signal. The presented algorithm offers to return both the filtered brain signal and the isolated artifact time course. The artifact signal can be used to assess the performance of the filter, as it should mainly exhibit the frequency characteristics of the stimulation signal. Analyses of the artifact time course could serve as a certainty proxy to assess current denoising performance, offering a potential reliability measure of the denoised brain activity over time.

The dynamic application of the spatial filter further reduces the amount of filtering by leaving all non-stimulation periods in the EEG signal unfiltered, thereby minimizing the overall signal manipulation to the necessary minimum. This prevents the filter being applied to and possibly distorting external noise during the stimulation-free intervals.

In conclusion, the presented algorithm features real-time, adaptive, and dynamically applied spatial filtering for the removal of EEG artifacts stemming from electrical neurostimulation, such as taVNS. The algorithm is designed to remain unaffected by alterations in the stimulation frequency and is capable of adapting to changes in the artifact topography, such as those resulting from impedance variations in the recording electrodes. This approach facilitates the development of closed-loop stimulation protocols with dynamic changes in stimulation parameters.

## 5 CRediT authorship contribution statement

David Menrath: Conceptualization and design, Methodology, Investigation, Software, Formal analysis, Writing – original draft. Joshua Woller: Conceptualization and design, Methodology, Investigation, Software, Formal analysis, Writing – original draft. Alireza Gharabaghi: Funding acquisition, Conceptualization and design, Project administration and supervision, Writing – review & editing.

This manuscript underwent careful revision by all authors and was further refined with regard to grammatical accuracy, sentence structure and wording using a convolutional neural network (DeepL) and an advanced language model (ChatGPT). We also used an Overleaf L^A^TEX template by George Kour to prepare the manuscript.

## 6 Data and code availability

Data and code supporting this study, as well as custom implementations of the denoising algorithms, will be made available by the first authors upon journal publication.

## 7 Funding

This investigator-initiated study was supported by the German Federal Ministry of Education and Research (BMBF) and the Ministry of Science, Research and the Arts of the State of Baden-Württemberg (MWK) through a grant from the German Center for Mental Health (DZPG Tü1C) and by the BMBF though a BEVARES grant (13GW0570B). The funding had no impact on the study design, on the collection, analysis and interpretation of data, on the writing of the report or on the decision to submit the article for publication.

## 8 Acknowledgements

We acknowledge support by the Open Access Publishing Fund of the University of Tübingen.

## 9 Declarations of competing interests

A.G. was supported by research grants from the German Federal Ministry of Education and Research (BMBF), the European Union’s Joint Program for Neurodegenerative Disease Research (EU-JPND), Medtronic, Abbott, and Boston Scientific, all of which were unrelated to this work.

